# Deconvoluting Metabolomic Flux in Living Cells with Real-time ^13^C J-coupling-edited Proton High-Resolution Magic Angle Spinning (HRMAS) NMR

**DOI:** 10.1101/2025.09.07.674640

**Authors:** Rajshree Ghosh Biswas, Ella Zhang, Aidan Pavao, Lynn Bry, Leo L. Cheng

**Author notes:** Corresponding Author: Leo Cheng, PhD, and Lynn Bry, MD, PhD. Authors contributed equally (Rajshree Ghosh Biswas, Ella Zhang).

## Abstract

Carbon-13 (^13^C) NMR provides a powerful approach to non-invasively investigate complex cellular metabolism. However, limitations arise per ^13^C NMR’s reduced sensitivity as compared to proton (^1^H) NMR, even when using ^13^C tracer molecules to enrich signal, and particularly when investigating metabolic processes in low volume samples. To overcome these limitations, we developed a ^1^H NMR method that harnesses ^1^H NMR’s sensitivity with ^13^C NMR’s specificity by capturing ^13^C NMR information in the ^1^H spectrum via ^13^C-^1^H J-couplings (^1^H[^13^C-Jed]). This approach improves ^13^C signal detection 13.9-fold and incorporates additional ^1^H NMR information to investigate complex metabolism on a cellular scale. Here, we demonstrate application of ^1^H[^13^C- Jed] NMR to deconvolute complex cellular metabolism in the anaerobic pathogen *Clostridioides difficile*. Analyses enabled precise tracking of branching metabolic reactions under anoxic and reducing conditions over time in a 30µL volume, and improved predictions of the pathogen’s genome scale metabolism. ^1^H[^13^C-Jed] NMR provides methodological advancements to investigate dynamic cellular metabolism and its role in complex biologic processes.

---

Complete understanding of cellular biologic functions, pathophysiological states and avenues for targeted therapies requires capacity to monitor real-time metabolic processes in living organisms. Nuclear magnetic resonance (NMR) spectroscopy provides a powerful tool for non-invasive monitoring of molecular changes, and of metabolic fluxes and pathway interactions in intact organisms.

^1^H NMR provides a common metabolic approach per the ^1^H nucleus’ high natural abundance, gyromagnetic ratio, and enhanced sensitivity. ^1^H one pulse NMR and Nuclear Overhauser Enhancement SpectroscopY (NOESY) with water suppression are commonly used for metabolomic profiling^1^. However, ^1^H NMR’s narrow chemical shift (0-10ppm) and associated spectral overlaps that occur among target molecules, given ^1^H-^1^H scalar coupling and spectral congestion among protons in similar chemical environments, limits its use. These effects make it difficult to identify metabolites and define metabolic dynamics in complex systems. While techniques to improve ^1^H NMR’s resolution, including the use of stronger magnetic fields, pure- shift NMR to remove signal multiplicity^2^, and multidimensional NMR (^1^H-^1^H COSY, TOCSY) to add additional signal, the high natural abundance of ^1^Hs in complex samples limits its capacity to effectively harness the rich information from ^1^H NMR in tracking complex metabolic dynamics in living organisms.

In contrast, use of defined ^13^C labelled substrates, with carbon’s larger spectral dispersion (0-200ppm) enables capacity to probe target metabolic pathways by tracking ^13^C-labelled metabolites in real-time. However, particularly in volume-limited samples, ^13^C NMR sensitivity remains well below that of ^1^H NMR and requires longer measurement times, rendering it difficult to observe fast reactions or control overall experiment times. NOE from ^1^H decoupling can enhance ^13^C sensitivity^3,4^, though the improvement remains modest relative to ^1^H NMR^5^. Recently, ^13^C labeled hyperpolarization has enabled improved sensitivity for monitoring fast metabolic processes *in vivo*^6–8^. However, ^13^C-hyperpolarized substrates have short relaxation times, on the order of seconds to minutes (e.g., ^13^C-1 pyruvate T^1^ relaxation is 2-3 mins)^9–11^, and only support monitoring of metabolic processes during this timeframe.

To address these issues, we developed a method that utilizes the sensitivity of ^1^H NMR to detect covalently bound ^13^C nuclei via the J-coupling that occurs between these NMR-active nuclei. To further advance the method, we use a scheme similar to heteronuclear single quantum coherence (HSQC) experiments that edits the ^1^H spectra through the lens of ^1^H-^13^C J-coupling, or ^1^H[^13^C-Jed]. When used for real-time quantification of metabolic flux in complex samples containing living, metabolically active cells, this approach produces a clean and high-resolution spectrum showing ^1^Hs bound to ^13^C.

We applied our approach to study dynamic metabolism in the obligate anaerobe *Clostridioides difficile.* This pathogen is a leading cause of healthcare acquired infections (HAIs) and causes substantive morbidity, mortality, and economic costs globally^13, 15, 26-28^. *C. difficile* has among the most diverse metabolic capabilities for human-infecting pathogens^13, 15^. It adapts its growth promoting metabolism within complex nutrient conditions by coordinating fermentations of amino acids, carbohydrates, short chain fatty acids, and even reductive metabolism of CO2 in the Wood-Ljungdahl pathway to produce 2-3 carbon backbones for anabolic processes^26-28^. Defining its metabolic strategies, both directly measured, and inferred from genome scale metabolic modeling, enables better approaches to prevent and treat these infections.

We conducted High-Resolution Magic Angle Spinning (HRMAS) NMR^12^ of metabolically active *C. difficile* in an experimental volume of 30µL, an approach that densely samples complex metabolism within a single reaction tube to enable robust inference of genome scale metabolism. ^1^H[^13^C-Jed] NMR provided superior and more sensitive information for genome scale metabolic flux over a 48 hour period^13–15^ than direct ^13^C NMR. ^1^H[^13^C-Jed] NMR enables analyses of living cells to identify metabolic contributions to our understanding of pathophysiologic processes and to identify targets for new therapeutic interventions.

## Results

We applied ^1^H[^13^C-Jed] to 100,000 *C. difficile* cells grown with uniformly labelled (U-^13^C) valine (**Figure 1a**), with and without ^13^C-decoupling, to show the spectral differences when using only ^13^C-labeled valine and products, compared to a standard 1D NOESY which detects all metabolites (Fig. 1b). In the latter case, the crowded ^1^H spectrum makes it difficult to evaluate metabolic products from metabolism of the U-^13^C valine, while ^1^H[^13^C-Jed] NMR enabled efficient monitoring of products and estimations of flux through their respective pathways. Decoupling also reduces spectral complexity, as can be seen in Fig.1, between decoupled (Fig. 1-i) and coupled acquisitions (Fig. 1-ii), both of which are comparable to the ^1^H[^13^C-Jed] and NOESY sensitivity of detection.

**Figure 1.**
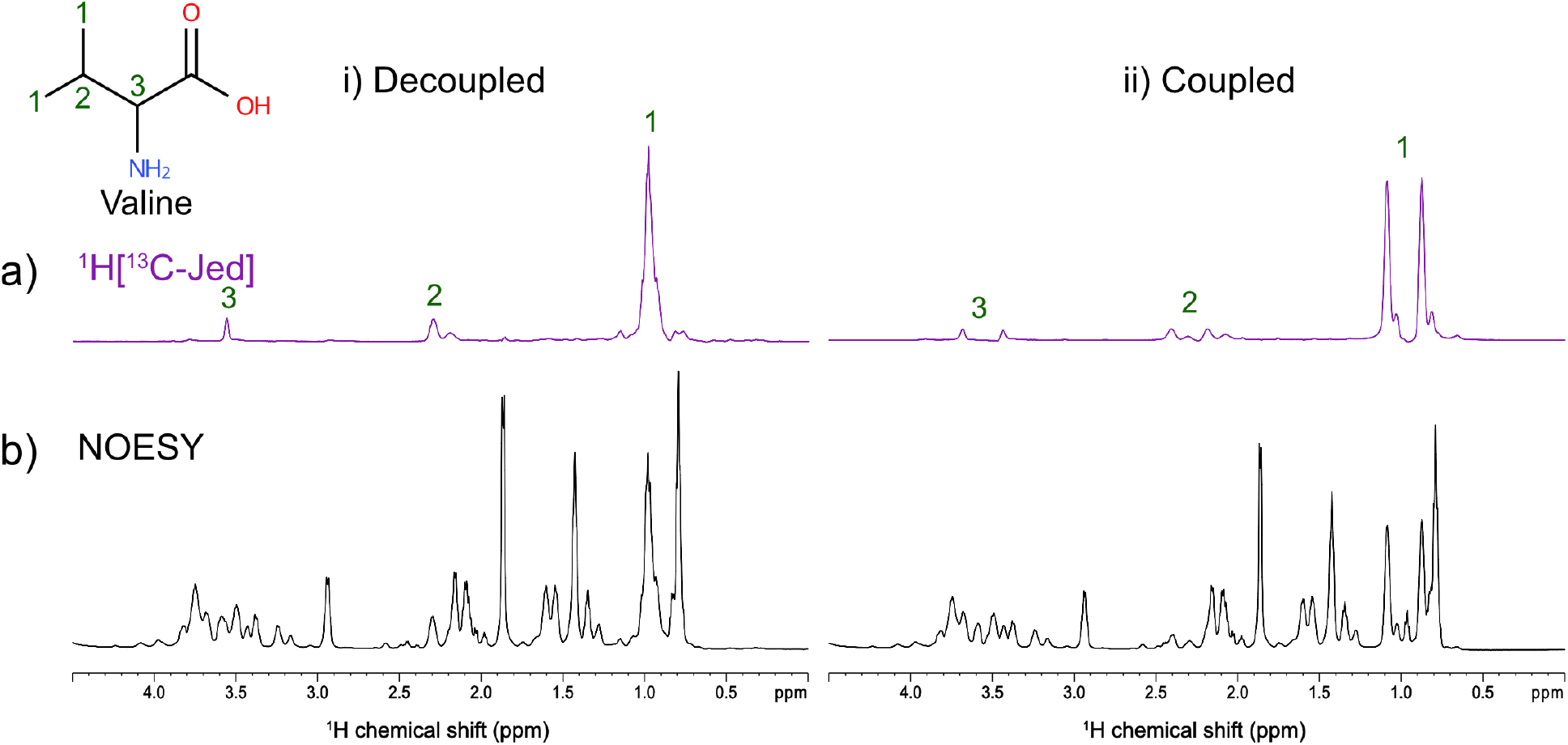
1D ^1^H *in vivo* HR-MAS NMR of *C. difficile* grown with U-^13^C valine comparing (a) ^1^H[^13^C-Jed] and (b) standard ^1^H NOESY with (i) and without (ii) ^13^C decoupling.

### ^13^C Glucose

We next applied ^1^H[^13^C-Jed] NMR to the more complex metabolism of uniformly labeled [U-^13^C]-glucose. As shown in **Figure 2,** anaerobic fermentation of [U-^13^C]-glucose to acetate, alanine, ethanol, and lactic acid by *C. difficile* over a 60-hour period can be clearly resolved^13^ with ^1^H[^13^C-Jed] NMR. Furthermore, acquisition of each ^1^H[^13^C-Jed] spectrum takes one third of the experimental time, at 12.68 mins for ^1^H[^13^C-Jed] versus 38.14 mins for ^13^C NMR, enabling higher resolution interrogation of rapid metabolic processes. The average signal to noise ratio (S/N) of glucose’s ^1^Hs attached to C-1 and carbon C-1, in ^1^H[^13^C-Jed] was 201±58 relative to direct ^13^C NMR which had an average S/N of 80±23. This improvement increased S/N ∼2.5x relative to ^13^C NMR, while also being 3x faster, with a time-savings factor of 18.75x. Note that the average S/N was calculated from the initial/starting experimental time point (at 0.84 hrs in ^13^C NMR and at 1.05 hrs in ^1^H[^13^C-Jed]), to avoid S/N discrepancies arising from changes due to microbial metabolism. The large standard deviation (201±58 vs 80±23) of the ^1^H[^13^C-Jed] S/N calculated from the S/N of the ^1^Hs attached to the C-1 of glucose may thus be owed to the increased sensitivity and reduced chemical shift range of ^1^H.

**Figure 2.**
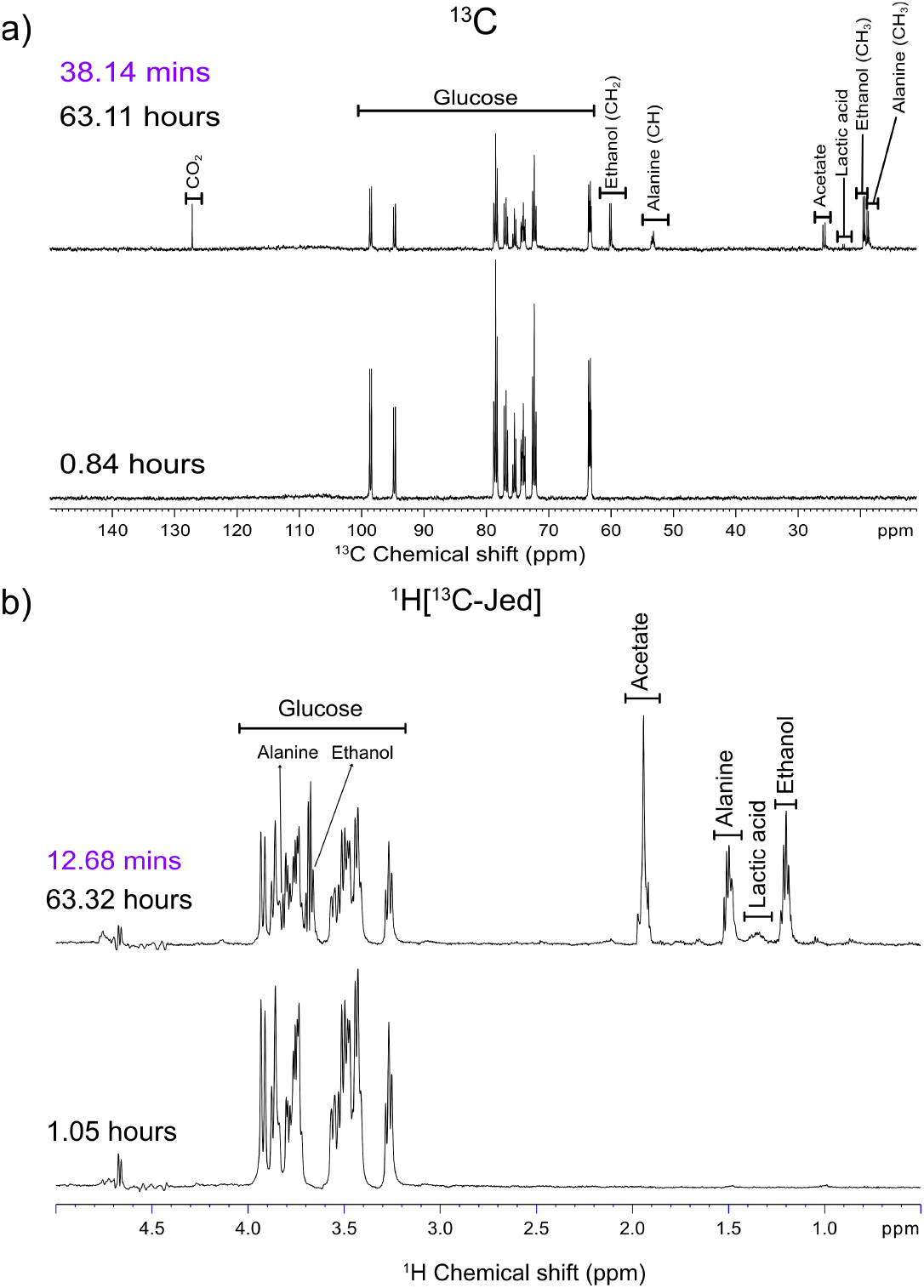
Monitoring live *C. difficile* cells supplemented with U- ^13^C glucose for over 63 hours using 1D HRMAS NMR. All metabolic products of glucose are labelled. a) Direct ^13^C NMR at 0.84 hours and 63.11 hours. B) Indirect ^1^H[^13^C-Jed] NMR at 1.05 hours and 63.32 hours. Each direct ^13^C experiment took 38.14 mins (purple) to collect compared to 12.68 mins (purple) of the indirect ^1^H[^13^C-Jed] NMR experiments.

Graphical depictions of U-^13^C glucose flux (**Figure 3**) showed comparable reaction rates for direct ^13^C and ^1^H[^13^C-Jed] NMR (Supplemental Table S1). The direct ^13^C NMR corresponds to a standard one-pulse, power gated carbon experiment with a 30° flip angle, while application of a ^1^H-decoupling regime during the relaxation delay utilizes the NOE from protons, thereby increasing signal intensity. However, when comparing Fig. 3a and 3b, the ^13^C NMR curve of alanine demonstrates higher intensity than that of ^1^H[^13^C-Jed] NMR. This is due to the use of alanine’s CH^3^ resonance for integration (Fig. 3a). When compared to the curve that used alanine’s CH moiety (Fig. 3c) the ^13^C and ^1^H[^13^C-Jed] NMR results more closely resemble each other. This finding illustrates an additional benefit of ^1^H[^13^C-Jed] NMR as compared to direct ^13^C NMR where NOE may impact signal intensity relative to, 1) the number of ^1^Hs attached to the carbon, and 2) the mobility of terminal CH_3_ groups versus less mobile CH moieties in the backbone structure^16,17^, thus impacting ^1^H-^13^C dipolar interactions and the longitudinal relaxation. This effect can also be seen in the ethanol curve (Fig. 3a vs 3b, green trajectory), where the direct ^13^C NMR measures the CH_2_ signal relative to one proton in the ^1^H[^13^C-Jed] experiment.

**Figure 3.**
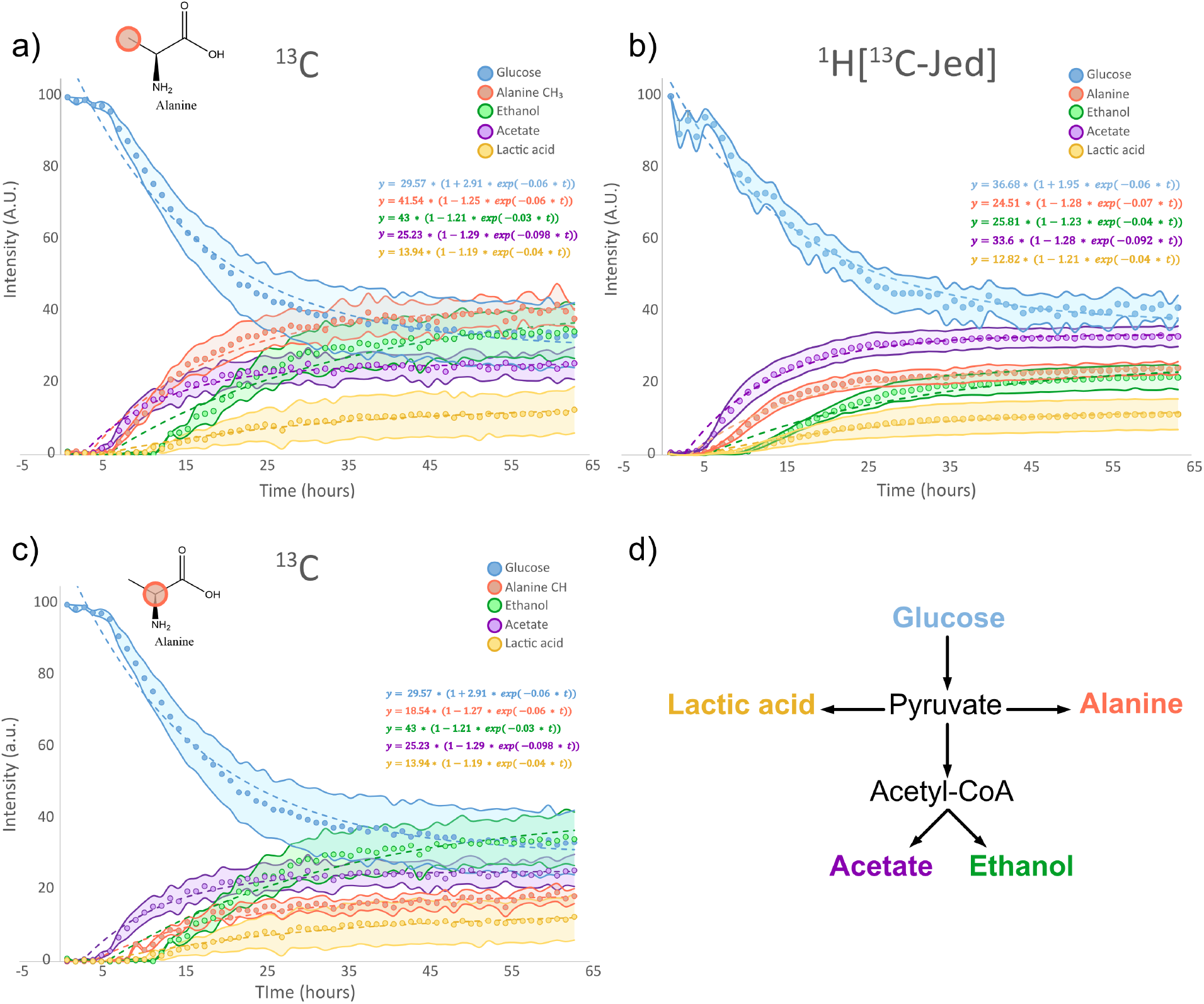
Graphical depiction of the metabolism of U-^13^C glucose (blue) to alanine (pink), ethanol (green), acetate (purple) and lactic acid (yellow) as a function of time (hours). Shaded error bounds represent standard errors from the triplicate studies. a) Direct ^13^C NMR where the CH_3_ doublet of alanine was integrated, b) indirect ^1^H[^13^C-Jed] NMR representing one proton of each metabolite and c) direct ^13^C NMR where the CH quartet of alanine was integrated to demonstrate that both direct ^13^C NMR and indirect ^1^H[^13^C-Jed] produce similar results. Here, the direct ^13^C NMR has the added consideration of influence from nuclear Overhauser enhancement (NOE) which increases the intensity of the alanine and ethanol graphs. D) schematic of the metabolism of glucose to the observed metabolic products. The intensities of the metabolic products were calculated by integrating the following regions: ^13^C glucose: 64-62.88ppm, ^13^C alanine CH_3_: 19.07-18.20ppm, ^13^C alanine CH: 53.8-52.5ppm, ^13^C ethanol: 60.7-59.4ppm, ^13^C acetate: 26.3- 25ppm, ^13^C lactic acid: 23-22.4 ppm. ^1^H glucose: 3.32-3.2 ppm, ^1^H alanine: 1.54 -1.44ppm, ^1^H ethanol: 1.25-1.15ppm, ^1^H acetate: 1.99-1.89ppm and ^1^H lactic acid: 1.42-1.29ppm.

### ^13^C Threonine

We next evaluated the performance of ^13^C and ^1^H[^13^C-Jed] NMR on *C. difficile* cells grown with U-^13^C threonine (Supporting figure S1), which the pathogen also ferments in branching oxidative and reductive pathways (**Figure 4**). Threonine’s analyses required additional improvements to the ^1^H[^13^C-Jed] method to resolve metabolic trajectories over time, including that its only distinct and measurable peak occurred close to water’s resonance at 4.2ppm.

**Figure 4.**
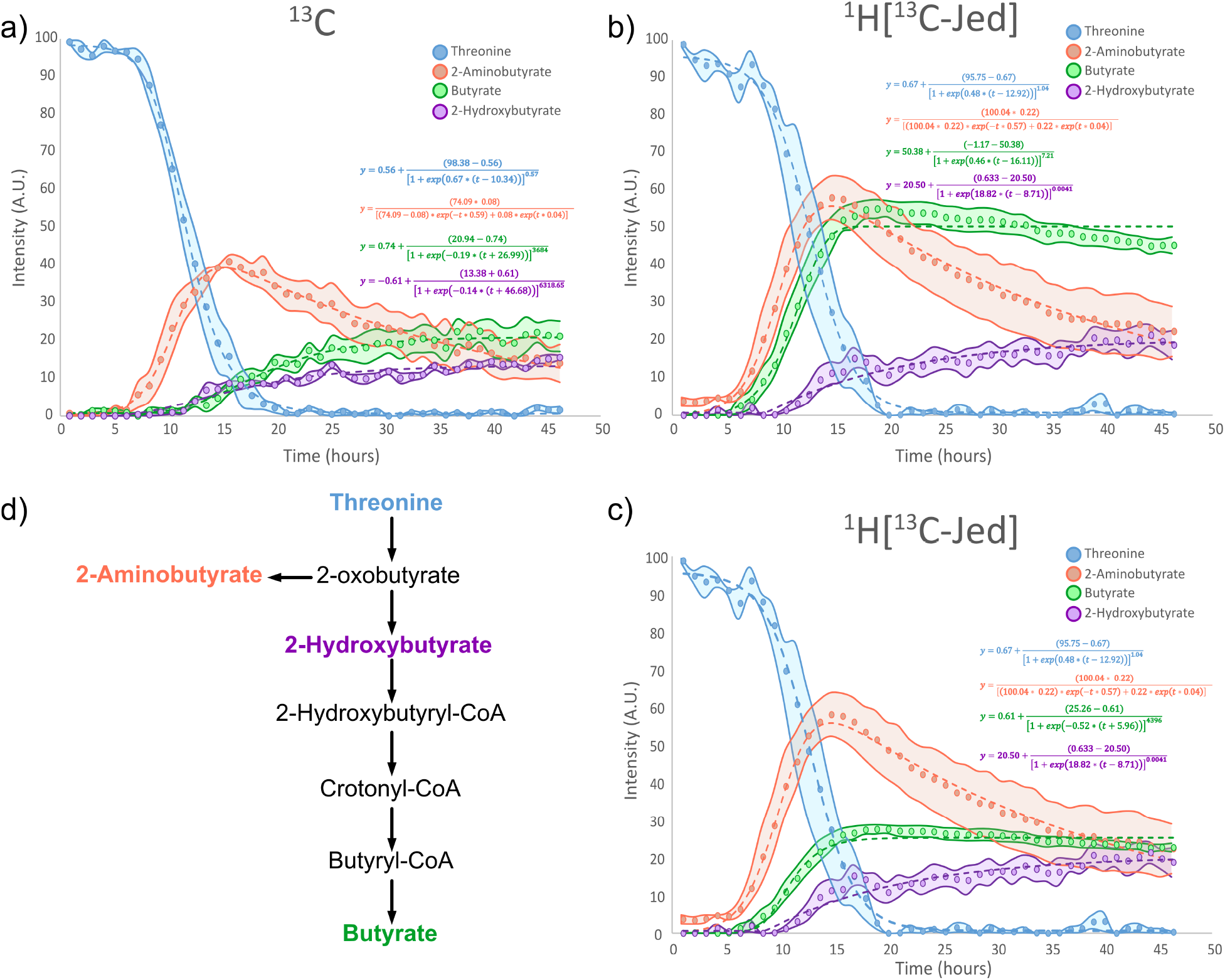
Graphical depiction of the metabolism of U-^13^C threonine (blue) to 2-aminobutyrate (pink), butyrate (green), and 2-hydroxybutyrate (purple) as a function of time (hours). Shaded error bounds represent standard errors from the triplicate studies. a) Direct ^13^C NMR, b) indirect ^1^H[^13^C-Jed] NMR representing one proton of each metabolite. Notice that the butyrate graph (green) is much larger in ^1^H[^13^C- Jed] compared to ^13^C NMR, due to spectral overlap resulting in the integration of propionate, and c) indirect ^1^H[^13^C-Jed] NMR where the 2 protons are of propionate are accounted for (divide the butyrate signal by 4) results in a graph that now closer resembles that of the direct ^13^C NMR. This demonstrates the considerations to ^1^H NMR, namely, due to the limited spectral dispersion of ^1^H NMR, many metabolic products can overlap making it difficult to follow individual metabolites. D) Schematic of the metabolism of threonine to the observed metabolic products. The intensities of the metabolic products were calculated by integrating the following regions: ^13^C threonine: 69.5-68.5 ppm, ^13^C 2-aminobutyrate: 26.81-25.95 ppm, ^13^C butyrate: 15.2-16.3 ppm, ^13^C 2-hydroxybutyrate: 30.19-29.12 ppm. ^1^H threonine: 4.3-4.17 ppm, ^1^H 2- aminobutyrate: 3.71-3.66 ppm, ^1^H butyrate: 2.22-2.08 ppm and, ^1^H 2-hydroxybutyrate: 4.03-3.91 ppm.

#### Water suppression

To avoid broad water resonances in ^1^H NMR, water suppression techniques are often used. However, due to the narrow spectral window of ^1^H NMR, any compound resonating near water’s resonance may also show suppression. This effect can be seen in the threonine curve (blue) in the ^1^H[^13^C-Jed] graph (Fig. 4b), where the curve has higher deviations at many time points when compared with the ^13^C curve. When possible, distinct signals away from the water should be observed in ^1^H[^13^C-Jed] NMR.

#### Spectral dispersion

^1^H NMR’s spectral dispersion over 0-10ppm is much less than that of ^13^C NMR (0-200ppm). When considering metabolic products with similar structures, including the starting substrate or other breakdown products, signal overlaps can make it difficult to discern and measure individual metabolites. This effect can be seen in Fig. 4b of the ^1^H[^13^C-Jed] spectrum (Fig. 4b), where the threonine-origin metabolites butyrate and propionate overlap in the ^1^H NMR at 2.1ppm. As such, the ^1^H[^13^C-Jed] curve for butyrate (green curve) was much larger than that of the ^13^C spectrum. However, if we consider the 2 ^1^Hs of propionate (Fig. 4c) the curve for butyrate more closely resembles that seen of its signatures in ^13^C NMR (Fig. 4a).

#### Spectral artifacts

The limited spectral dispersion of ^1^H NMR can also introduce spectral artifacts, such as ^1^H-^13^C J-coupling satellite signals that may mask metabolites of interest. This effect can be addressed by optimizing the decoupling regimes^18^. Moreover, a spinning speed of 3600 Hz was sufficient at moving spinning sidebands, particularly from water, spaced 6 ppm apart at 600 MHz, to outside of the ^1^H spectral window. We previously demonstrated that *C. difficile* cells readily withstand centripetal forces at a spinning rate of 5000 Hz^19^. However, when using a 4 mm HRMAS rotor, more fragile cells or tissues may require lower spinning rates at <3000 Hz to maintain sample integrity^20–22^. This change can introduce spinning sidebands which manifest as artifacts occurring as increments of the spinning speed, thus masking signals of interest. Techniques that avoid or remove spinning sidebands include optimizing water suppression under MAS^20,23^ as well as approaches to handle in post-processing of datasets^24^ or using variable spinning magic angle spinning (VSMAS)^25^ NMR.

### Optimizing Experimental Parameters for ^1^H[^13^C-Jed] NMR

When investigating live-cells in dense time series, experimental parameters require effective optimization to maximize the signal-to-noise (S/N). Here, we optimized variations in the recycle delay (RD) time where the S/N is represented as a unit of the measurement time for ^1^H[^13^C- Jed] and ^13^C HRMAS NMR. We performed these analyses with a 20µL solution of 150mM 1,2- ^13^C ornithine (**Figure 5**). By varying the RD time, we calculated the S/N per unit time for the ^13^C- 2 and the ^1^H attached to the ^13^C-2.

**Figure 5.**
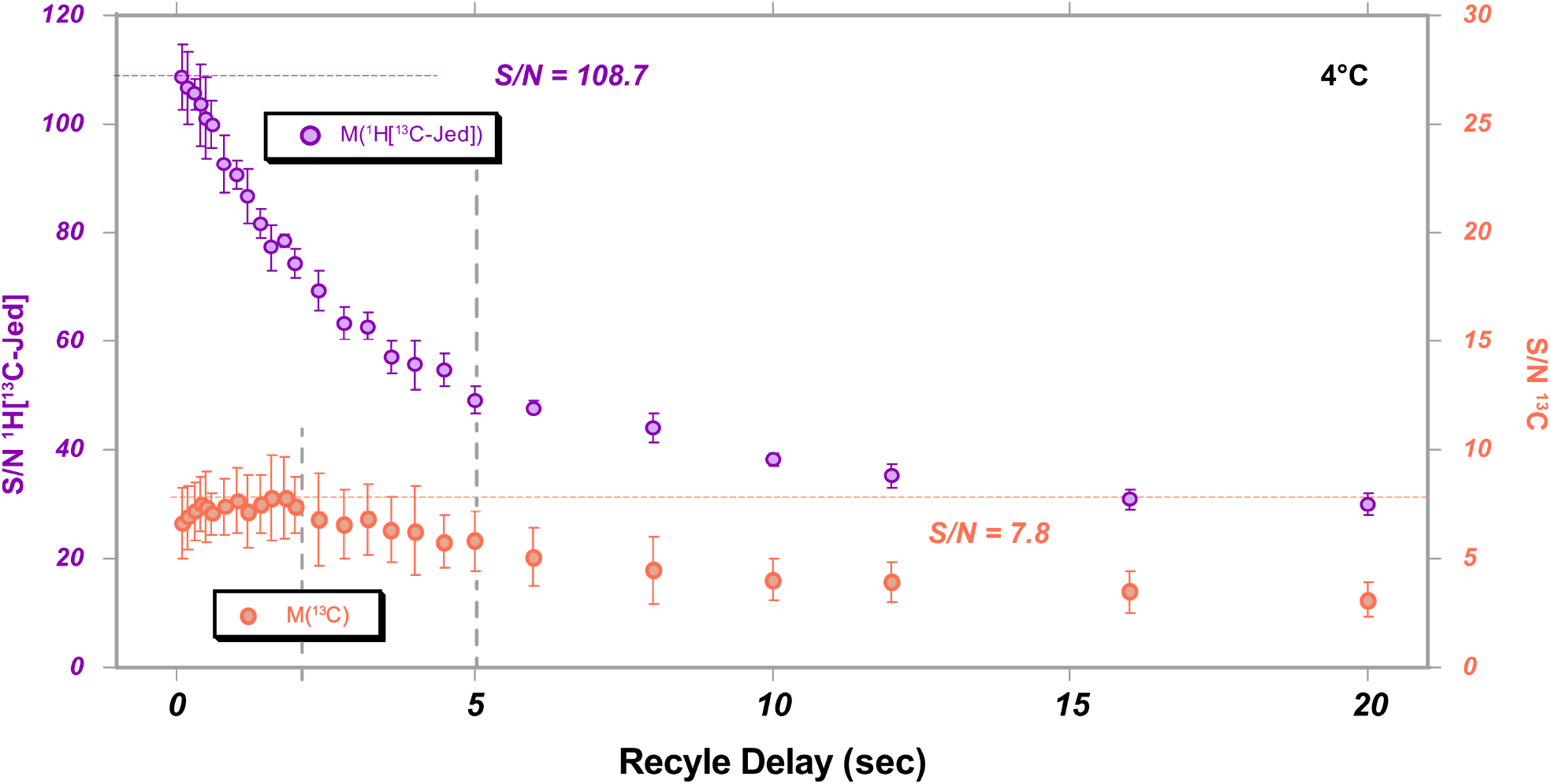
Optimization of the recycle delay (RD in seconds) during direct ^13^C NMR (pink) and indirect ^1^H[^13^C-Jed] NMR (purple) for max signal to noise (S/N) using 150 mM 1,2-^13^C ornithine. Here the ^13^C-2 carbon and the ^1^H directly attached to ^13^C-2 were measured. The max S/N of direct ^13^C NMR was calculated to be 7.8, whereas that of ^1^H[^13^C-Jed] was 108.7. This corresponds to a time saving factor of 194 if labelled carbons were indirectly observed by detecting the attached protons. The grey dashed lines represent the recycle delays used for the previous experiments (glucose and threonine) without optimization, where ^13^C NMR (RD of 2 s) was optimal; however, in the ^1^H[^13^C-Jed] NMR experiments (RD of 5 s), approximately half the optimal S/N was measured. With a shorter recycle delay, more scans can be collected with faster experiment times using ^1^H[^13^C-Jed] NMR. The caveat here is that the results are not quantitative and bias the shorter relaxing (T_1_) functional groups of the metabolites.

As shown in Fig. 5, analyses identified a maximum S/N of 108.7 for ^1^H[^13^C-Jed] when using a RD of 1s, and a S/N of 7.8 for ^13^C NMR when using a RD of 2s. These findings provided a 13.9-fold improvement in S/N for ^1^H[^13^C-Jed] NMR relative to ^13^C NMR, corresponding to a 194-fold reduction in experimental time for the tested RDs.

In the absence of optimization, these approaches nonetheless provided substantive improvements in S/N and reductions in experimental time for the analyses shown in Figures 3 and 4, with an 8.3-fold improvement in S/N of ^1^H[^13^C-Jed] NMR versus ^13^C NMR, or 69-fold reduction in experimental time (Fig 5; vertical dotted grey line depicts the RD=5s for ^1^H[^13^C-Jed]).

## Discussion

We present application of ^1^H[^13^C-Jed] NMR to advance real-time investigation of *C. difficile’s* anaerobic metabolism. This approach greatly improved the sensitivity of detection of substrates and metabolites, with substantive reductions in sampling time to enable denser interrogation of complex and rapid biochemical reactions. *C. difficile’s* capacity to concurrently metabolize disparate substrates enables its pathogenesis within the gut^26–28,29^ where it must compete with other commensal microbes, and with the host for ever changing pools of nutrients. High-resolution understanding of the pathogen’s metabolism is thus essential to identify and develop improved preventive and therapeutic strategies. This approach also provides broader opportunities to study additional pathogens, symbionts, and species of economic or environmental importance.

We demonstrate the superiority of ^1^H[^13^C-Jed] HRMAS NMR to delineate complex metabolomic flux for *C. difficile.* Analyses need only a limited input of 100,000 cells within a 30µL reaction volume to support continuous monitoring for 48 hours or longer. By observing ^1^Hs bound to ^13^C nuclei we leverage the inherent sensitivity of ^1^H over ^13^C NMR, while avoiding common problems with congested resonances that plague traditional 1D ^1^H experiments. This approach improves the detection of metabolites from input substrates and estimation of their concentrations^13^. When leveraged as input into genome scale metabolic models, the improved information enables broader inference of cellular metabolism to define target pathways, substrates and cellular systems for further investigation by NMR and other methods^13^.

1D direct ^13^C and ^13^C-edited ^1^H NMR has been used to elucidate energy metabolism^5^ and neurotransmission^30^ by following metabolic flux in rat^31–33^ and human brains^34,35^. With the sensitivity of ^1^H in addition to the enhanced specificity achieved via ^13^C labeling, mass/volume limited samples (e.g., cells) can be observed in their native state. It should be noted that proton-observed carbon editing (POCE, ^1^H[^13^Ced]), first introduced by Rothman et al.,^34^ uses both a non-selective spin-echo alongside a ^13^C inversion pulse on alternating scans, to further separate ^1^Hs attached and not attached to ^13^C, i.e., to differentiate protons bound to ^12^C vs ^13^C.^35^

Our studies using uniformly ^13^C enriched substrates did not use the selective ^13^C inversion pulse, thus enabling observation of all ^1^Hs directly attached to ^13^C, thereby excluding signal from natural abundance substrates in the media, where the frequency of ^13^C is approximately 1% for any given carbon. By using ^1^H[^13^C-Jed] HRMAS NMR, the ^1^H spectrum specifically detects the ^13^C-labelled substrates and associated metabolic products.

A consideration of direct ^13^C NMR is the influence of NOE to enhance the sensitivity of low gyromagnetic nuclei in volume-limited samples. When comparing the metabolism of U-^13^C glucose by *C. difficile* (Fig. 3), we observed differences in sensitivity versus direct ^13^C and ^1^H[^13^C- Jed] NMR, particularly in the evolution of alanine and ethanol. By choosing a different carbon to integrate, the graph more closely resembled that of ^1^H[^13^C-Jed], thus demonstrating the variability in sensitivity when using NOE-derived experiments. To avoid the influence of NOE, an inverse gated ^1^H-decoupling one-pulse carbon experiment (ZGIG, where decoupling is only on during the acquisition) can be used. However, when considering the small sample volume for *in vivo* microbial studies, any sensitivity enhancement is advantageous.

We also acknowledge considerations associated with traditional 1D ^1^H NMR including. 1) need to optimize methods used for water suppression. 2) Effects of limited spectral dispersion, where metabolic by-products with similar structures can overlap, but can be alleviated through the indirect observations of covalently bonded ^13^C nuclei and inform the need to monitor distinct metabolite peaks as possible. 3) Spectral artifacts, such as spinning sidebands in HRMAS NMR, which can often be addressed by optimizing spinning rates to reduce effects while preserving sample integrity. 4) Susceptibility distortion in gel-like samples, such as cell cultures, which can be alleviated by using HRMAS to narrow the broadened line shapes that arise from chemical shift anisotropy (CSA) and dipolar interactions. However, when working with time-sensitive reactions, samples must often be loaded and evaluated immediately to capture rapid metabolic processes. In such cases, it’s critical to first optimize acquisition parameters, shimming and field homogeneity, else ^1^H line shapes become susceptible to distortions that inaccurately measure metabolite concentrations.

With optimization for the system under study, ^1^H[^13^C-Jed] provides major advancements to observe complex metabolism of ^13^C-labelled substrates, in a manner that cannot be done with standard 1D ^1^H or ^13^C NMR alone. The approach has benefits over hyperpolarization given the capacity for lengthy, dense time series that greatly exceed the limited relaxation time of hyperpolarized substrates. As such ^1^H[^13^C-Jed] NMR can provide complementary information of metabolic processes that extend over hours to days.

Our approach further supports high-resolution, longitudinal analyses of volume limited samples (<30µL), and can include the use of smaller HRMAS inserts (e.g., 12µL spherical inserts)^21^, thereby improving the filling factor and overall sensitivity of detection of ^13^C substrates and metabolites. The reduced sample diameter of inserts further reduces centripetal forces on samples, enabling the analysis of delicate cells and tissues. In addition, magic angle coil spinning (MACS)^36,37^, where a tuned micro-coil surrounds and spins with the sample and inductively couples with the outer detection coil, can increase the sensitivity of NMR to nano-liter levels on intact samples.^38^ These techniques, with optimized recycle delay (Fig. 5) at the Ernst angle, or flip angle that maximizes signal intensity, enables fast reaction monitoring with increased sensitivity. These methods can be used singly, and in combination, to identify target metabolic pathways for therapeutic interventions leveraging alternative substrate analogues or other biotherapeutics such as commensal microorganisms with more efficient and competing pathways to consume pathogen- preferred substrates^39^.

While the study of anaerobe metabolism has long been more difficult than that of aerotolerant species, the sealed HRMAS inserts enable high-resolution investigations. In contrast, studies of obligate aerobes or other aerotolerant species to evaluate responses in atmospheric O_2_, require access to oxygen for the duration of the experiment^40^. For such cases, studies by Simpson, *et al.* devised HR-MAS rotor caps with ∼500 µm holes drilled to permit oxygen exchange, enabling *in vivo* monitoring of metabolic processes in freshwater shrimp.^41^ Their work demonstrated that the centripetal force under MAS creates a vortex that brings in oxygen, reducing anoxic stress and increasing the longevity of the organism^41^.

^1^H[^13^C-Jed] NMR may also be advantageous for targeted, solution-state process monitoring using low-field NMR, which is inherently limited by its spectral resolution^42,43. 1^H[^13^C- Jed] NMR’s versatility holds potential to monitor longitudinal dynamics for a variety of processes including metabolic fluxes in intact cells and whole organisms, under normal or disease-promoting conditions^34,44,45^, environmental, physiochemical and toxicity responses in living organisms^41,46,47^. The approach also supports high-resolution studies of enzyme kinetics^48,49^, electrochemical reactions^50^ and more. ^1^H[^13^C-Jed] NMR provides a simple, efficient and highly sensitive technique to observe complex dynamics in metabolic and other chemical reactions, especially in mass/volume-limited samples.

## Material and Methods

### C. difficile cells

*C. difficile* Pathogenicity Locus-deleted (PaLoc-) ATCC 43255 was cultured as described in Pavao et al.^13^ in Modified Minimal Media (MM) containing 100 µM sodium selenite and the respective uniformly labelled substrates: 27.5mM ^13^C-Glucose, 15mM ^13^C- Threonine or 15mM ^13^C-Valine. Following culturing, cells were diluted to 100,000 CFU and prepared into a 30 µL HRMAS insert along with 2 µL of D_2_O (spectrometer lock) and sealed. This insert was then placed into a 4 mm zirconia rotor for NMR analysis.

### NMR spectroscopy

Experiments were performed on a Bruker Avance III 600 MHz (^1^H) spectrometer equipped with a 4 mm HRMAS ^1^H-^13^C-^2^H (lock) probe fitted with a magic angle gradient. Samples were spun at 3600 Hz under MAS. All experiments were locked on D_2_O and maintained at 37° C. A pre-saturation water suppression technique was applied to all ^1^H NMR experiments. The metabolism of the labelled substrates by *C. difficile* cells were monitored continuously (for >48 hours) using alternating 1D:

1. ^1^H Nuclear Overhauser Enhancement Spectroscopy (NOESY) collected with a mixing time of 200ms, 16K time-domain points, 20ppm sweep width, 5 s recycle delay. All experiments were collected with 4 dummy scans and 128 total scans. Spectra were processed on Topspin 3.6.2 (Bruker Biospin) with manual phase and baseline corrections, and 0.3 Hz line broadening.
2. ^13^C one-pulse power-gated experiment with a 30° flip angle collected with 16K time-domain points, 240 ppm sweep width, and 2 s recycle delay. A low power WALTZ16 ^1^H decoupling regime was used during both the recycle delay and acquisition. All experiments were collected using 4 dummy scans and 1024 total scans. Spectra were processed on Topspin 3.6.2 (Bruker BioSpin) with manual phase and baseline corrections, and 5 Hz line broadening.
3. One dimensional heteronuclear correlation spectroscopy (HSQC) resulting in ^1^H[^13^C-Jed] NMR. This information was collected in phase sensitive mode with states-TPPI. Here, the F2 dimension was collected using 8K time-domain points and 1 increment in the F1 dimension. All experiments were collected using 24 ppm spectral width, 5 s recycle delay, 16 dummy scans and 128 total scans. A low power GARP-4 decoupling regime was used. All spectra were processed on Topspin 3.6.2 (Bruker Biospin) with manual phase and baseline corrections, and 0.3 Hz line broadening.

Non-linear curve fit models and equations were determined using JMP Pro 16 (SAS Institute, Cary, NC, USA).

### Ornithine

A 150 mM solution of 1,2-^13^C ornithine was prepared in Dulbecco’s Phosphate Buffer Saline (dPBS) and D_2_O and inserted into a 50 µL HRMAS insert and sealed. The sample was then placed into a 4 mm zirconia NMR rotor for analysis. The sample was spun at 5000 Hz, locked on D_2_O and maintained at 25° C.

The carbon experiments with a 30° flip angle were collected with 64K time-domain points, a spectral width of 240 ppm, 4 dummy scans and 32 total scans.

The proton experiments (^1^H[^13^C-Jed]) were collected with 32K time domain points, a spectral width of 24 ppm, a recycle delay of 16 dummy scans, and 32 total scans.

For optimization, the recycle delay was varied from 0.2 - 20s (0.2, 0.3, 0.4, 0.5, 0.6, 0.8, 1, 1.2, 1.4, 1.6, 1.8, 2, 2.4, 2.8, 3.2, 3.6, 4, 4.5, 5, 6, 8, 10, 12, 14, 16, 18, 20s). The signal to noise ratio of ^13^C-2 and the ^1^H attached to the ^13^C-2 were calculated and plotted (Fig. 5: S/N vs recycle delay).

## Acknowledgements

This project was supported by the National Institutes of Health (L.B., grant nos. R01AI153653, R03AI174158 and P30DK056338; L.L.C., grant nos. S10OD023406, R21CA243255 and R01AG070257), the Brigham and Women’s Hospital Precision Medicine Institute and Presidential Scholar’s Award (L.B.), the MGH A. A. Martinos Center for Biomedical Imaging (L.B. and L.L.C.) and the Massachusetts Life Sciences Center (L.B. and L.L.C.). We would also like to thank the Canadian Institute of Health Research (CIHR) for the following postdoctoral training fellowship: MFE-194064 (R.G.B).

## Author Contributions

L.B and L.L.C conceived the study. A.P and L.B developed the anaerobic microbiology methods and performed all microbiological aspects of the experiments. L.L.C developed the HRMAS NMR methodologies and E.Z and L.L.C performed the magnetic resonance spectroscopy components of the HRMAS experiments. R.G.B and L.L.C analyzed the NMR data. R.G.B and L.L.C wrote the original draft. L.B provided edits and content to the original draft.

## Data availability

Data is available from the authors upon request.

## References

(1) Nagana Gowda, G. A.; Raftery, D. Can NMR Solve Some Significant Challenges in Metabolomics? J. Magn. Reson. 2015, 260, 144–160. 10.1016/j.jmr.2015.07.014.

(2) Zangger, K. Pure Shift NMR. Prog. Nucl. Magn. Reson. Spectrosc. 2015, 86–87, 1–20. 10.1016/j.pnmrs.2015.02.002.

(3) Sailasuta, N.; Robertson, L. W.; Harris, K. C.; Gropman, A. L.; Allen, P. S.; Ross, B. D. Clinical NOE 13C MRS for Neuropsychiatric Disorders of the Frontal Lobe. J. Magn. Reson. 2008, 195 (2), 219–225. 10.1016/j.jmr.2008.09.012.

(4) Mandal, P. K.; Guha Roy, R.; Samkaria, A.; Maroon, J. C.; Arora, Y. In Vivo 13C Magnetic Resonance Spectroscopy for Assessing Brain Biochemistry in Health and Disease. Neurochem. Res. 2022, 47 (5), 1183–1201. 10.1007/s11064-022-03538-8.

(5) De Graaf, R. A.; Mason, G. F.; Patel, A. B.; Behar, K. L.; Rothman, D. L. In Vivo 1 H-[13 C]-NMR Spectroscopy of Cerebral Metabolism. NMR Biomed. 2003, 16 (6–7), 339–357. 10.1002/nbm.847.

(6) Sharma, G.; Enriquez, J. S.; Armijo, R.; Wang, M.; Bhattacharya, P.; Pudakalakatti, S. Enhancing Cancer Diagnosis with Real-Time Feedback: Tumor Metabolism through Hyperpolarized 1-13C Pyruvate MRSI. Metabolites 2023, 13 (5), 606. 10.3390/metabo13050606.

(7) Keshari, K. R.; Kurhanewicz, J.; Jeffries, R. E.; Wilson, D. M.; Dewar, B. J.; Van Criekinge, M.; Zierhut, M.; Vigneron, D. B.; Macdonald, J. M. Hyperpolarized 13 C Spectroscopy and an NMR-compatible Bioreactor System for the Investigation of Real-time Cellular Metabolism. Magn. Reson. Med. 2010, 63 (2), 322–329. 10.1002/mrm.22225.

(8) Rider, O. J.; Apps, A.; Miller, J. J. J. J.; Lau, J. Y. C.; Lewis, A. J. M.; Peterzan, M. A.; Dodd, M. S.; Lau, A. Z.; Trumper, C.; Gallagher, F. A.; Grist, J. T.; Brindle, K. M.; Neubauer, S.; Tyler, D. J. Noninvasive In Vivo Assessment of Cardiac Metabolism in the Healthy and Diabetic Human Heart Using Hyperpolarized 13 C MRI. Circ. Res. 2020, 126 (6), 725–736. 10.1161/CIRCRESAHA.119.316260.

(9) Serrao, E. M.; Brindle, K. M. Potential Clinical Roles for Metabolic Imaging with Hyperpolarized [1-13C]Pyruvate. Front. Oncol. 2016, 6. 10.3389/fonc.2016.00059.

(10) Pagès, G.; Tan, Y. L.; Kuchel, P. W. Hyperpolarized [1, 13 C]Pyruvate in Lysed Human Erythrocytes: Effects of Co-substrate Supply on Reaction Time Courses. NMR Biomed. 2014, 27 (10), 1203–1210. 10.1002/nbm.3176.

(11) Brindle, K. M. Imaging Metabolism with Hyperpolarized 13 C-Labeled Cell Substrates. J. Am. Chem. Soc. 2015, 137 (20), 6418–6427. 10.1021/jacs.5b03300.

(12) Cheng, L. L. High-resolution Magic Angle Spinning NMR for Intact Biological Specimen Analysis: Initial Discovery, Recent Developments, and Future Directions. NMR Biomed. 2023, 36 (4), e4684. 10.1002/nbm.4684.

(13) Pavao, A.; Girinathan, B.; Peltier, J.; Altamirano Silva, P.; Dupuy, B.; Muti, I. H.; Malloy, C.; Cheng, L. L.; Bry, L. Elucidating Dynamic Anaerobe Metabolism with HRMAS 13C NMR and Genome-Scale Modeling. Nat. Chem. Biol. 2023, 19 (5), 556–564. 10.1038/s41589-023-01275-9.

(14) Pavao, A.; Zhang, E.; Monestier, A.; Peltier, J.; Dupuy, B.; Cheng, L.; Bry, L. HRMAS 13 C NMR and Genome-Scale Metabolic Modeling Identify Threonine as a Preferred Dual Redox Substrate for Clostridioides Difficile. September 18, 2023. 10.1101/2023.09.18.558167.

(15) Pavao, A.; Girinathan, B.; Peltier, J.; Silva, P. A.; Dupuy, B.; Cheng, L. L.; Bry, L. Real-Time HRMAS 13 C NMR of Obligately Anaerobic Cells Identifies New Metabolic Targets in the Pathogen Clostridioides Difficile. May 4, 2021. 10.1101/2021.05.04.442336.

(16) Macdonald, P. M.; Soong, R. The Truncated Driven NOE and 13C NMR Sensitivity Enhancement in Magnetically-Aligned Bicelles. J. Magn. Reson. 2007, 188 (1), 1–9. 10.1016/j.jmr.2007.06.002.

(17) Dvinskikh, S. V.; Yamamoto, K.; Dürr, U. H. N.; Ramamoorthy, A. Sensitivity and Resolution Enhancement in Solid-State NMR Spectroscopy of Bicelles. J. Magn. Reson. 2007, 184 (2), 228–235. 10.1016/j.jmr.2006.10.004.

(18) Bahadoor, A.; Brinkmann, A.; Melanson, J. E. 13 C-Satellite Decoupling Strategies for Improving Accuracy in Quantitative Nuclear Magnetic Resonance. Anal. Chem. 2021, 93 (2), 851–858. 10.1021/acs.analchem.0c03428.

(19) Vedantam, G.; Clark, A.; Chu, M.; McQuade, R.; Mallozzi, M.; Viswanathan, V. K. Clostridium Difficile Infection: Toxins and Non-Toxin Virulence Factors, and Their Contributions to Disease Establishment and Host Response. Gut Microbes 2012, 3 (2), 121–134. 10.4161/gmic.19399.

(20) Taylor, J. L.; Wu, C. L.; Cory, D.; Gonzalez, R. G.; Bielecki, A.; Cheng, L. L. High-Resolution Magic Angle Spinning Proton NMR Analysis of Human Prostate Tissue with Slow Spinning Rates. Magn. Reson. Med. 2003, 50 (3), 627–632. 10.1002/MRM.10562.

(21) Aime, S.; Bruno, E.; Cabella, C.; Colombatto, S.; Digilio, G.; Mainero, V. HR-MAS of Cells: A “Cellular Water Shift” Due to Water-protein Interactions? Magn. Reson. Med. 2005, 54 (6), 1547–1552. 10.1002/mrm.20707.

(22) André, M.; Dumez, J.-N.; Rezig, L.; Shintu, L.; Piotto, M.; Caldarelli, S. Complete Protocol for Slow-Spinning High-Resolution Magic-Angle Spinning NMR Analysis of Fragile Tissues. Anal. Chem. 2014, 86 (21), 10749–10754. 10.1021/ac502792u.

(23) Mobarhan, Y. L.; Struppe, J.; Fortier-McGill, B.; Simpson, A. J. Effective Combined Water and Sideband Suppression for Low-Speed Tissue and in Vivo MAS NMR. Anal. Bioanal. Chem. 2017, 409 (21), 5043–5055. 10.1007/s00216-017-0450-3.

(24) Burns, M. A.; Taylor, J. L.; Wu, C.-L.; Zepeda, A. G.; Bielecki, A.; Cory, D.; Cheng, L. L. Reduction of Spinning Sidebands in Proton NMR of Human Prostate Tissue with Slow High-Resolution Magic Angle Spinning. Magn. Reson. Med. 2005, 54 (1), 34–42. 10.1002/mrm.20523.

(25) Madhu, P. K.; Pratima, R.; Kumar, A. Suppression of Sidebands by Variable Speed Magic Angle Sample Spinning in Solid State NMR. Chem. Phys. Lett. 1996, 256 (1–2), 87–89. 10.1016/0009-2614(96)00428-9.

(26) Girinathan, B. P.; DiBenedetto, N.; Worley, J. N.; Peltier, J.; Arrieta-Ortiz, M. L.; Immanuel, S. R. C.; Lavin, R.; Delaney, M. L.; Cummins, C. K.; Hoffman, M.; Luo, Y.; Gonzalez-Escalona, N.; Allard, M.; Onderdonk, A. B.; Gerber, G. K.; Sonenshein, A. L.; Baliga, N. S.; Dupuy, B.; Bry, L. In Vivo Commensal Control of Clostridioides Difficile Virulence. Cell Host Microbe 2021, 29 (11), 1693-1708.e7. 10.1016/j.chom.2021.09.007.

(27) Arrieta-Ortiz, M. L.; Immanuel, S. R. C.; Turkarslan, S.; Wu, W.-J.; Girinathan, B. P.; Worley, J. N.; DiBenedetto, N.; Soutourina, O.; Peltier, J.; Dupuy, B.; Bry, L.; Baliga, N. S. Predictive Regulatory and Metabolic Network Models for Systems Analysis of Clostridioides Difficile. Cell Host Microbe 2021, 29 (11), 1709-1723.e5. 10.1016/j.chom.2021.09.008.

(28) Allegretti, J. R.; Marcus, J.; Storm, M.; Sitko, J.; Kennedy, K.; Gerber, G. K.; Bry, L. Clinical Predictors of Recurrence After Primary Clostridioides Difficile Infection: A Prospective Cohort Study. Dig. Dis. Sci. 2020, 65 (6), 1761–1766. 10.1007/s10620-019-05900-3.

(29) Smits, W. K.; Lyras, D.; Lacy, D. B.; Wilcox, M. H.; Kuijper, E. J. Clostridium Difficile Infection. Nat. Rev. Dis. Primer 2016, 2 (1), 16020. 10.1038/nrdp.2016.20.

(30) Rothman, D. L.; De Graaf, R. A.; Hyder, F.; Mason, G. F.; Behar, K. L.; De Feyter, H. M. In Vivo 13 C and 1 H-[13 C] MRS Studies of Neuroenergetics and Neurotransmitter Cycling, Applications to Neurological and Psychiatric Disease and Brain Cancer. NMR Biomed. 2019, 32 (10), e4172. 10.1002/nbm.4172.

(31) Rothman, D. L.; Behar, K. L.; Hetherington, H. P.; Den Hollander, J. A.; Bendall, M. R.; Petroff, O. A.; Shulman, R. G. 1H-Observe/13C-Decouple Spectroscopic Measurements of Lactate and Glutamate in the Rat Brain in Vivo. Proc. Natl. Acad. Sci. 1985, 82 (6), 1633–1637. 10.1073/pnas.82.6.1633.

(32) De Graaf, R. A.; Brown, P. B.; Mason, G. F.; Rothman, D. L.; Behar, K. L. Detection of [1,6-13 C 2]-glucose Metabolism in Rat Brain by in Vivo 1 H-[13 C]-NMR Spectroscopy. Magn. Reson. Med. 2003, 49 (1), 37–46. 10.1002/mrm.10348.

(33) De Graaf, R. A.; Chowdhury, G. M. I.; Behar, K. L. Quantification of High-Resolution 1 H-[13 C] NMR Spectra from Rat Brain Extracts. Anal. Chem. 2014, 86 (10), 5032–5038. 10.1021/ac5006926.

(34) Rothman, D. L.; Novotny, E. J.; Shulman, G. I.; Howseman, A. M.; Petroff, O. A.; Mason, G.; Nixon, T.; Hanstock, C. C.; Prichard, J. W.; Shulman, R. G. 1H-[13C] NMR Measurements of [4-13C]Glutamate Turnover in Human Brain. Proc. Natl. Acad. Sci. 1992, 89 (20), 9603–9606. 10.1073/pnas.89.20.9603.

(35) Chen, H.; De Feyter, H. M.; Brown, P. B.; Rothman, D. L.; Cai, S.; De Graaf, R. A. Comparison of Direct 13 C and Indirect 1 H-[13 C] MR Detection Methods for the Study of Dynamic Metabolic Turnover in the Human Brain. J. Magn. Reson. 2017, 283, 33–44. 10.1016/j.jmr.2017.08.004.

(36) Sakellariou, D.; Goff, G. L.; Jacquinot, J.-F. High-Resolution, High-Sensitivity NMR of Nanolitre Anisotropic Samples by Coil Spinning. Nature 2007, 447 (7145), 694–697. 10.1038/nature05897.

(37) Lucas-Torres, C.; Wong, A. Current Developments in µMAS NMR Analysis for Metabolomics. Metabolites 2019, 9 (2), 29. 10.3390/metabo9020029.

(38) Adhikari, S. S.; Zhao, L.; Dickmeis, T.; Korvink, J. G.; Badilita, V. Inductively Coupled Magic Angle Spinning Microresonators Benchmarked for High-Resolution Single Embryo Metabolomic Profiling. Analyst 2019, 144 (24), 7192–7199. 10.1039/C9AN01634A.

(39) Bae, J.; Park, K.; Kim, Y.-M. Commensal Microbiota and Cancer Immunotherapy: Harnessing Commensal Bacteria for Cancer Therapy. Immune Netw. 2022, 22 (1), e3. 10.4110/in.2022.22.e3.

(40) Judge, M. T.; Wu, Y.; Tayyari, F.; Hattori, A.; Glushka, J.; Ito, T.; Arnold, J.; Edison, A. S. Continuous in Vivo Metabolism by NMR. Front. Mol. Biosci. 2019, 6, 26. 10.3389/fmolb.2019.00026.

(41) Mobarhan, Y. L.; Fortier-McGill, B.; Soong, R.; Maas, W. E.; Fey, M.; Monette, M.; Stronks, H. J.; Schmidt, S.; Heumann, H.; Norwood, W.; Simpson, A. J. Comprehensive Multiphase NMR Applied to a Living Organism. Chem. Sci. 2016, 7 (8), 4856–4866. 10.1039/c6sc00329j.

(42) Kreyenschulte, D.; Paciok, E.; Regestein, L.; Blümich, B.; Büchs, J. Online Monitoring of Fermentation Processes via Non-invasive Low-field NMR. Biotechnol. Bioeng. 2015, 112 (9), 1810– 1821. 10.1002/bit.25599.

(43) Jenne, A.; Soong, R.; Downey, K.; Biswas, R. G.; Decker, V.; Busse, F.; Goerling, B.; Haber, A.; Simpson, M. J.; Simpson, A. J. Brewing Alcohol 101: An Undergraduate Experiment Utilizing Benchtop NMR for Quantification and Process Monitoring. Magn. Reson. Chem. 2024, 62 (6), 429– 438. 10.1002/mrc.5428.

(44) Peng, Y.; Zhang, Z.; He, L.; Li, C.; Liu, M. NMR Spectroscopy for Metabolomics in the Living System: Recent Progress and Future Challenges. Anal. Bioanal. Chem. 2024, 416 (9), 2319–2334. 10.1007/s00216-024-05137-8.

(45) Li, Z.; Zheng, L.; Shi, J.; Zhang, G.; Lu, L.; Zhu, L.; Zhang, J.; Liu, Z. Toxic Markers of Matrine Determined Using 1 H-NMR-Based Metabolomics in Cultured Cells In Vitro and Rats In Vivo. Evid. Based Complement. Alternat. Med. 2015, 2015, 1–11. 10.1155/2015/598412.

(46) Gomis, M. I.; Vestergren, R.; Borg, D.; Cousins, I. T. Comparing the Toxic Potency in Vivo of Long-Chain Perfluoroalkyl Acids and Fluorinated Alternatives. Environ. Int. 2018, 113, 1–9. 10.1016/j.envint.2018.01.011.

(47) Nami, F.; Ferraz, M. J.; Bakkum, T.; Aerts, J. M. F. G.; Pandit, A. Real-Time NMR Recording of Fermentation and Lipid Metabolism Processes in Live Microalgae Cells. Angew. Chem. 2022, 134 (14), e202117521. 10.1002/ange.202117521.

(48) Roy, A. D.; Jayalakshmi, K.; Dasgupta, S.; Roy, R.; Mukhopadhyay, B. Real Time HR–MAS NMR: Application in Reaction Optimization, Mechanism Elucidation and Kinetic Analysis for Heterogeneous Reagent Catalyzed Small Molecule Chemistry. Magn. Reson. Chem. 2008, 46 (12), 1119–1126. 10.1002/mrc.2321.

(49) Anderson, L. A.; Franz, A. K. Real-Time Monitoring of Transesterification by 1 H NMR Spectroscopy: Catalyst Comparison and Improved Calculation for Biodiesel Conversion. Energy Fuels 2012, 26 (10), 6404–6410. 10.1021/ef301035s.

(50) Albert, K.; Dreher, E.; Straub, H.; Rieker, A. Monitoring Electrochemical Reactions by 13 C NMR Spectroscopy. Magn. Reson. Chem. 1987, 25 (10), 919–922. 10.1002/mrc.1260251017.

