## Supplemental Data for "Deconvoluting Metabolomic Flux in Living Cells with Real-time ^13^C J-coupling-edited Proton High-Resolution Magic Angle Spinning (HRMAS) NMR"

ǂ Authors contributed equally

* Corresponding Author:

Leo Cheng, PhD

### Section 1: Glucose metabolism by *C. difficile in vivo*

**Table S1:** $^{13}C$ and $^{1}H[^{13}C\text{-Jed}]$ Reaction rates of the metabolism of U-$^{13}C$ Glucose to Alanine, Ethanol, Acetate and Lactic acid as depicted in the graphs within Figure 3 in the main text. There are two $^{13}C$ graphs for the production alanine where either the $CH_3$ doublet (Fig. 3A) or the CH quartet (Fig. 3C) are integrated. As they are the same metabolite, the reaction rate of alanine does not change, suggesting that the changes in intensity between the two groups are attributed to the NOE.

| **Metabolite** | **$^{13}C$ NMR** | **$^{1}H[^{13}C\text{-Jed}]$** |
|---|---|---|
| Glucose | 0.06 | 0.06 |
| Alanine ($CH_3$) | 0.06 | 0.07 |
| Alanine (CH) | 0.06 | |
| Ethanol | 0.03 | 0.04 |
| Acetate | 0.098 | 0.092 |
| Lactic acid | 0.04 | 0.04 |

### Section 2: Threonine metabolism by *C. difficile in vivo*

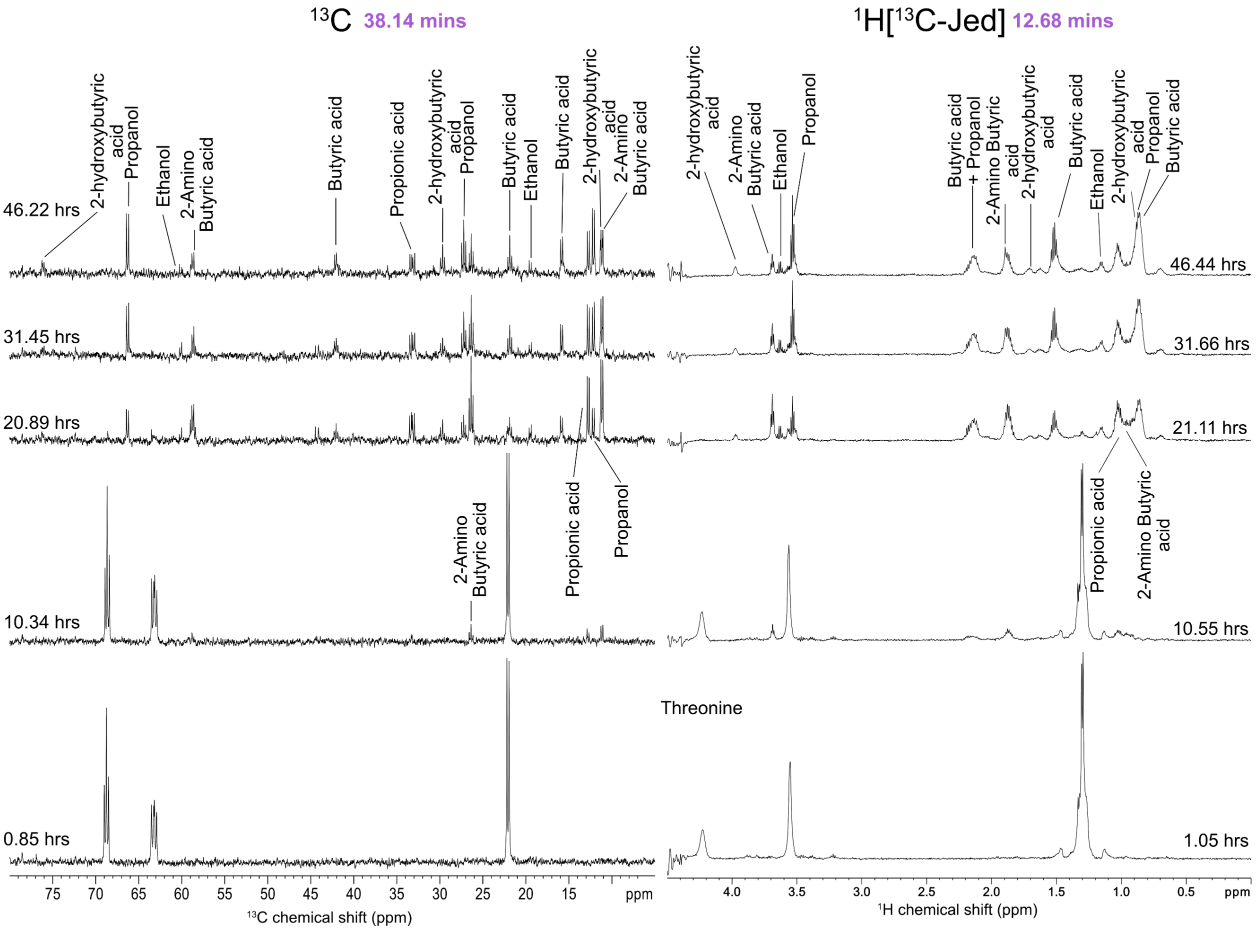


**Figure S1:** Direct $^{13}C$ and indirect $^{1}H[^{13}C\text{-Jed}]$ NMR monitoring the metabolism of U-$^{13}C$ threonine by *C. difficile* cells, *in vivo,* at 5 time points for over 46 hours. All metabolic products of threonine are labelled. Note that at each timepoint, the $^{13}C$ spectra took 38.14 mins to collect compared to that of $^{1}H[^{13}C\text{-Jed}]$ NMR which only took 12.68 mins. Here, the average signal to noise ratio (SNR, across triplicate studies) of threonine for the $^{1}Hs$ attached to C-2 and carbon C-2, in $^{1}H[^{13}C\text{-Jed}]$ was $113 \pm 20$ relative to direct $^{13}C$ NMR which had an average SNR of $20 \pm 3$. $^{1}H[^{13}C\text{-Jed}]$ exhibits 5.6x increase in SNR relative to $^{13}C$ NMR, while also being 3x faster, corresponding to a time-saving factor of 32x.

Table S2: $^{13}C$ and $^{1}H[^{13}C\text{-Jed}]$ Reaction rates of the metabolism of U-$^{13}C$ Threonine to 2-Aminobutyrate, Butyrate and 2-Hydroxybutyrate as depicted in the graphs within Figure 4 in the main text. Butyrate (i) represents the $^{1}H[^{13}C\text{-Jed}]$ reaction rate of butyrate including the overlapping propionate resonance (Fig. 4b), whereas butyrate (ii) represents the $^{1}H[^{13}C\text{-Jed}]$ reaction rate of butyrate when accounting for the 2 additional protons of propionate. The reaction rates will differ between $^{13}C$ NMR and $^{1}H[^{13}C\text{-Jed}]$ due to differences in production time & rate of propionate which overlaps with the butyrate signal. The difference in the reaction rate of 2-hydroxybutyrate may be attributed to the influence of water suppression in $^{1}H[^{13}C\text{-Jed}]$ NMR as well as the reduced sensitivity of $^{13}C$ NMR. This is a consideration of $^{1}H[^{13}C\text{-Jed}]$ NMR resulting from the limited spectral dispersion of $^{1}H$ nuclei.

| **Metabolite** | **$^{13}C$ NMR** | **$^{1}H[^{13}C\text{-Jed}]$** |
|---|---|---|
| Threonine | 0.67 | 0.48 |
| 2-Aminobutyrate | -0.59 | -0.57 |
| Butyrate (i) | -0.19 | 0.46 |
| Butyrate (ii) | | -0.52 |
| 2-Hydroxybutyrate | -0.14 | 18.82 |
